# Flexible Path Planning Through Vicarious Trial and Error

**DOI:** 10.1101/2021.09.08.459317

**Authors:** Jeffrey L. Krichmar, Nicholas A. Ketz, Praveen K. Pilly, Andrea Soltoggio

**Affiliations:** Department of Cognitive Sciences, Department of Computer Science, University of California, Irvine, Irvine, CA 92697-5100; Center for Human-Machine Collaboration, Information and Systems Sciences Laboratory, HRL Laboratories, 3011 Malibu Canyon Road, Malibu, CA 90265, USA; Computer Science Department, Loughborough University, Loughborough LE11 3TU, UK

**Keywords:** Cognitive map, Hippocampus, Navigation, Preplay, Spiking Neural Network

## Abstract

Flexible planning is necessary for reaching goals and adapting when conditions change. We introduce a biologically plausible path planning model that learns its environment, rapidly adapts to change, and plans efficient routes to goals. Unlike prior models of hippocampal replay, our model addresses the decision-making process when faced with uncertainty. We tested the model in simulations of human and rodent navigation in mazes. Like the human and rat, the model was able to generate novel shortcuts, and take detours when familiar routes were blocked. Similar to rodent hippocampus recordings, the neural activity of the model resembles neural correlates of Vicarious Trial and Error (VTE) during early learning or during uncertain conditions. Similar to rodent studies, after learning, the neural activity resembles forward replay or preplay predicting a future route, and VTE activity decreases. We suggest that VTE, in addition to weighing possible outcomes, is a way in which an organism may gather information for future use.

## 1 Introduction

Flexible planning is an important aspect of cognition that is especially useful when achieving a goal under uncertain conditions. Multi-step planning can be thought of as a process that uses a cognitive map to guide a sequence of actions towards a goal [Miller and Venditto, 2021]. In the context of spatial navigation, humans and other animals have the ability to choose alternate routes when necessary. Moreover, they can express spatial knowledge in the form of novel shortcuts over locations not previously explored [Chrastil and Warren, 2014, Boone et al., 2019].

Neural correlates of planning have been observed in the dynamic responses of place cells in the rodent hippocampus. In hippocampal replay, plans are formed by reactivating place cells of previously experienced locations. In hippocampal preplay, place cells activate in sequence according to future trajectories that may be taken by the animal [Dragoi and Tonegawa, 2011, Pfeiffer and Foster, 2013]. Computational models have shown how hippocampal preplay can plan towards new goals in familiar environments and replan when familiar routes are no longer viable [Mattar and Daw, 2018, Stachenfeld et al., 2017]. However, models of replay do not address the decision-making process when faced with uncertainty, and they may not explain how knowledge of never experienced locations can be stored and expressed.

In the mid-twentieth century, Edward Tolman described the flexible, intelligent behavior observed in animals as a cognitive map [Tolman, 1948]. One aspect of a cognitive map was Vicarious Trial and Error (VTE), which is the ability to weigh one’s options before taking decisive action. Similar to hippocampal preplay, neural correlates of VTE have been observed in hippocampal CA1 where neurons with place-specific firing exhibit “sweeps” of activity representing the locations at which the animal looks while considering its left versus right turn choice [Johnson and Redish, 2007, Redish, 2016]. VTE seems to occur when the animal is uncertain about which route to take. After experience, hippocampal preplay occurs for the path the animal intends to take, but not the alternative routes.

In this paper, we introduce a computational model of path planning that demonstrates VTE during early learning or when environmental conditions change. We further show that this model can acquire knowledge about never experienced paths, and rapidly adapt to express this knowledge when challenged. To simulate VTE and path planning, we used a spiking wavefront propagation path planner [Hwu et al., 2018]. The spiking wavefront propagation path planner is like other diffusion-based planners used in robotics [Choset et al., 2005]. However, the present model has features that make it more neurobiologically plausible. Neurons in the spiking neural network represent place cells and preplay can be observed in the activity of the neurons during the planning stage. In Hwu et al. [2018], the map of the environment was given *a priori*. In the present paper, the environmental features, such as wall, boundaries and other obstacles, are initially unknown. To learn the map, we extended the spiking wavefront propagation model with a biologically plausible learning rule called E-Prop [Bellec et al., 2020]. E-Prop was designed to learn sequences in spiking recurrent neural networks. In the present paper, E-Prop is used to learn efficient navigation routes to goals.

We demonstrate flexible path planning and correlates of VTE by comparing the model to human navigation experiments, a detour task used in rodents, and in an open field task (i.e., Morris water maze). We show that neural population activity is greater when the simulated agent is uncertain about which way to go, due to unfamiliar environments or unexpected changes to the environment. We suggest this activity is comparable to VTE observed in animals. Similar to rodent studies [Redish, 2016], once the environment becomes familiar, neural population activity more resembles hippocampal preplay. Our results suggest that VTE may assist in the acquisition of knowledge that can be later expressed as novel shortcuts or rapid re-routing.

This work provides two major contributions:

- We extend the spiking wavefront propagation path planner [Hwu et al., 2018] to learn the map of the environment through E-Prop [Bellec et al., 2020].
- We demonstrate a model of VTE like behavior in three environments and compare to both human and rodent behavioral data.

## 2 Methods

### 2.1 Spiking Wave Propagation

Spiking wavefront propagation is a neuromorphic navigation algorithm inspired by neuronal dynamics and connectivity of neurons in the brain [Hwu et al., 2018]. The algorithm is loosely based on the dynamic responses of place cells in the hippocampus. In hippocampal preplay, place cells activate in sequence according to future trajectories that may be taken by the animal [Dragoi and Tonegawa, 2011, Pfeiffer and Foster, 2013]. Spiking wavefront propagation is also supported by the observation that spreading activation of brain activity is seen in several areas of the brain including the hippocampus [Zhang and Jacobs, 2015].

The spiking wavefront propagation algorithm learns by adjusting axonal delays. This was inspired by biological evidence suggesting that the myelin sheath, which wraps around and insulates axons, may undergo a form of activity-dependent plasticity [Fields, 2015]. These studies have shown that the myelin sheath becomes thicker with learning motor skills and cognitive tasks. A thicker myelin sheath implies faster conduction velocities and improved synchrony between neurons. Adjusting axonal propagation fits better with the idea behind wave propagation than the more commonly used synaptic weight updates, which change the ability of a synapse to contribute to the firing of a post-synaptic neuron, but not necessarily the timing of propagating an action potential.

The spiking wavefront propagation algorithm assumed a grid representation of space. Each grid unit corresponded to a discretized area of physical space, and connections between units represented the ability to travel from one grid location to a neighboring location. Each unit in the grid represents a single neuron with spiking dynamics, which are captured with a model described by equations below, which have also been described in [Hwu et al., 2018].

The membrane potential of neuron *i* at time *t* + 1 is represented by Eqn 1:

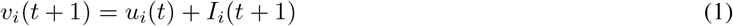

in which *u*_*i*_(*t*) is the recovery variable and *I*_*i*_(*t*) is the input current at time t.

The recovery variable *u*_*i*_(*t*+) is described Eqn 2:

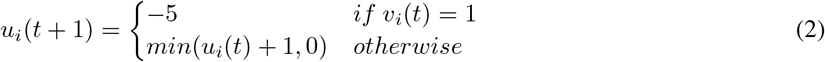

such that immediately after a membrane potential spike, the recovery variable starts as a negative value and linearly increases toward a baseline value of 0.

The input current I at time t+1 is given by Eqn 3:

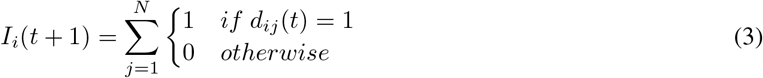

such that *d*_*ij*_(*t*) is the delay counter of the signal from neighboring neuron *j* to neuron *i*. This delay is given by Eqn 4:

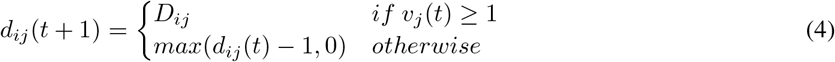

which behaves as a timer corresponding to an axonal delay with a starting value of *D*_*ij*_(*t*), counting down until it reaches 0.

The value of *D*_*ij*_(*t*) depends on the environmental cost associated with the spatial area between neurons *i* and *j*. Input currents come from neighboring connected neurons. When a neighboring neuron spikes, the input current *I*_*i*_ is increased by 1. This triggers the neuron to spike. After the spike, the recovery variable *u*_*i*_ is set to -5, then gradually recovers back to 0, modeling the refractory period. Also, immediately after spiking, all delay counters *d*_*ij*_ for all neighbor neurons *j* are set to their current values of *D*_*ij*_. In the present paper, and in contrast to Hwu et al. [2018], *D*_*ij*_ are initially set to some arbitrary value and then learned through exploration. These values are updated based on the agent’s observations in the environment with a learning rule described in the section on E-Prop and Backpropagation Through Time.

### 2.2 E-Prop and Back-Propagation Through Time

The E-Prop algorithm was introduced to learn sequences in spiking recurrent neural networks [Bellec et al., 2020]. Learning was dictated by a loss function related to the desired output. The credit assignment problem was resolved by subjecting each neuron to an eligibility trace based on the neuron’s recent activity. Weights between neurons were updated based on the loss and the value of the eligibility trace. In this way, the E-Prop algorithm resembled Back-Propagation Through Time (BPTT).

In the present model, E-Prop was used to learn sequences of movements through an environment based on the traversal cost. As discussed in the previous section, the spiking wavefront propagation algorithm calculates a path. The neurons activated during the wave propagation are eligible to be updated. Once eligible, they are subject to an exponential decay due to an eligibility trace. The most eligible neurons are those most recently active relative to when the wave reaches the goal destination. After the path is calculated, E-Prop is applied to weights projecting from neurons along the calculated path. Since these weights are connected to locations adjacent to the path, we assume the agent can observe the features (e.g., traversal cost) at these map locations. In this way, E-Prop solves the credit assignment problem by rewarding paths that lead to goal locations, while also learning about the environmental structure of nearby map locations.

In the spiking wavefront propagation neural network, weights denote the axonal delay in sending an action potential or spike from the pre-synaptic neuron to the post-synaptic neuron. We applied the E-Prop algorithm to these weights so that the spiking neural network would learn an axonal delay corresponding to environmental features observed in *map*_*xy*_, which is the spatial location at Cartesian coordinates (*x, y*) of the *map*.

The values of *D*_*ij*_ are updated when the agent reaches the goal destination. The learning rule is described by Eqn 5:

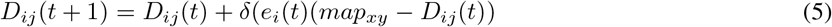

where *δ* is the learning rate, set to 0.5, *e*_*i*_(*t*) is an eligibility trace for neuron *i*, and *map*_*xy*_ represents the observed cost for traversing the location (*x, y*), which corresponds to neuron *i*. This rule is applied for each of the neighboring neurons, *j*, of neuron *i*. By using this method of axonal plasticity, the agent can simultaneously explore and learn, adapting to changes in the environment. The loss in Eqn. 5 is *map*_*xy*_ - *D*_*ij*_.

The eligibility trace for neuron *i* is given by Eqn 6:

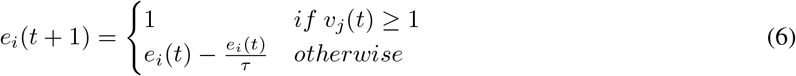

where *τ* is the rate of decay for the eligibility trace. The setting of *τ* will be explored in the next section.

### 2.3 Extracting a Path from the Spike Wavefront Algorithm

We illustrate the spiking wavefront algorithm with a simple example (see Figure 1). In this example, there is a 4×4 spiking neural network representing the space (see Figure 1A). The inner section of the of the environment contains an obstacle that is twice as costly to traverse than the outer section.

**Figure 1:**
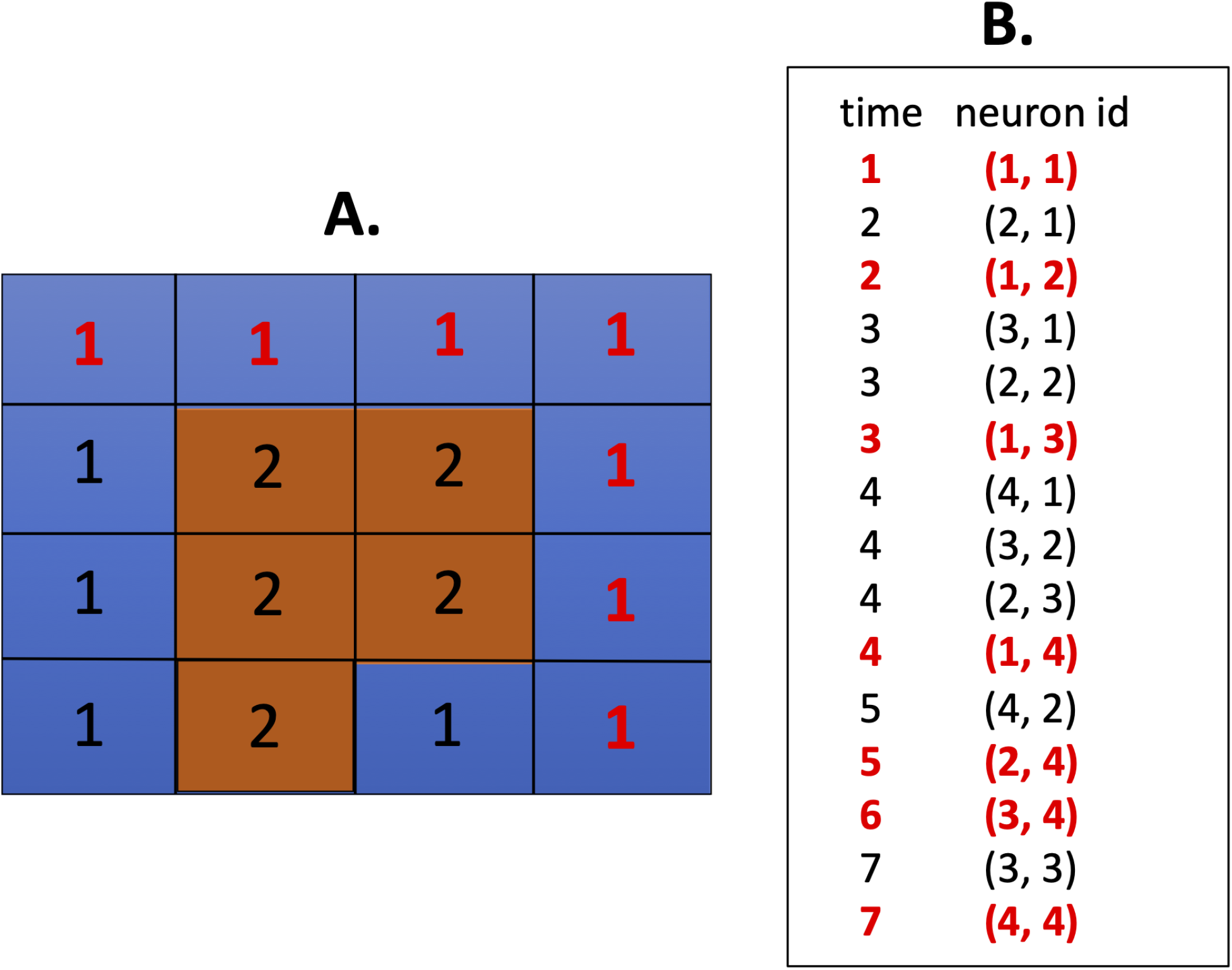
Simple example with a 4×4 neural network. A neuron is connected to its 4 neighbors (North, South, East, West). The task is to find a path from (1,1) to (4,4). The resulting path found by the spike wave propagation algorithm is denoted in red font. A. 4×4 neural network representing a simple environment. The delays, which correspond to a cost map of the environment, are numbered. B. Example of using the spike table for extracting a path from (1,1) to (4,4).

To use the path planning algorithm with delay-encoded costs, the neuron corresponding to the current location (1,1) is induced to spike. This causes an input current to be sent to neighboring neurons, starting a traveling wave of activity across the area covered by the grid. As each neuron spikes, the spike index and the time of spike are logged in a table. Figure 1B illustrates how this information is used to trace a path from start to goal location (4,4).

To extract a path from the spike table, a list of neuron IDs is maintained, starting with the goal neuron. The first spike time of the goal neuron is found (see 1B). Then, the timestamps are decremented until the spiking of a neuron neighboring the goal neuron is found. That neuron is then added to the list. The process then continues by finding spikes of neurons neighboring the most recent neuron. These are appended to the list until the start neuron is added. The result is the optimal path between the start and goal in reversed order. This method of extracting paths has similarities to evidence suggesting the nervous system decodes information based on spike time rankings [VanRullen et al., 2005].

## 3 Results

The spiking wave propagation algorithm with E-Prop learning was tested in three different environments: 1) Human navigation in a virtual maze [Boone et al., 2019], 2) Rodent navigation in the Tolman detour maze [Alvernhe et al., 2011], and 3) Rodent navigation in the Morris Water Maze [Morris, 1984]. We simulated their experimental protocols and compared the simulation results with the human and animal experiments.

### 3.1 Simulating Human Navigation and Taking Novel Shortcuts

In Boone et al. [2019], human participants navigated through a virtual hedge maze where they first took several laps on a fixed path, and then were tested by how well they could navigate between locations (see Figure 2). Along the route, there were landmarks, such as a chair, mailbox, potted plant, a picnic table, which participants were to take note of. During the test phase, participants were placed at one landmark and told to navigate toward another landmark. Some subjects took the learned path between landmarks, and others took novel shortcuts to landmarks. In another experiment, participants were told to take the shortest path to a landmark. This resulted in more participants taking novel shortcuts, and suggested that participants had survey knowledge, an allocentric mental map of the environment, even if they had previously not chosen to express this knowledge.

**Figure 2:**
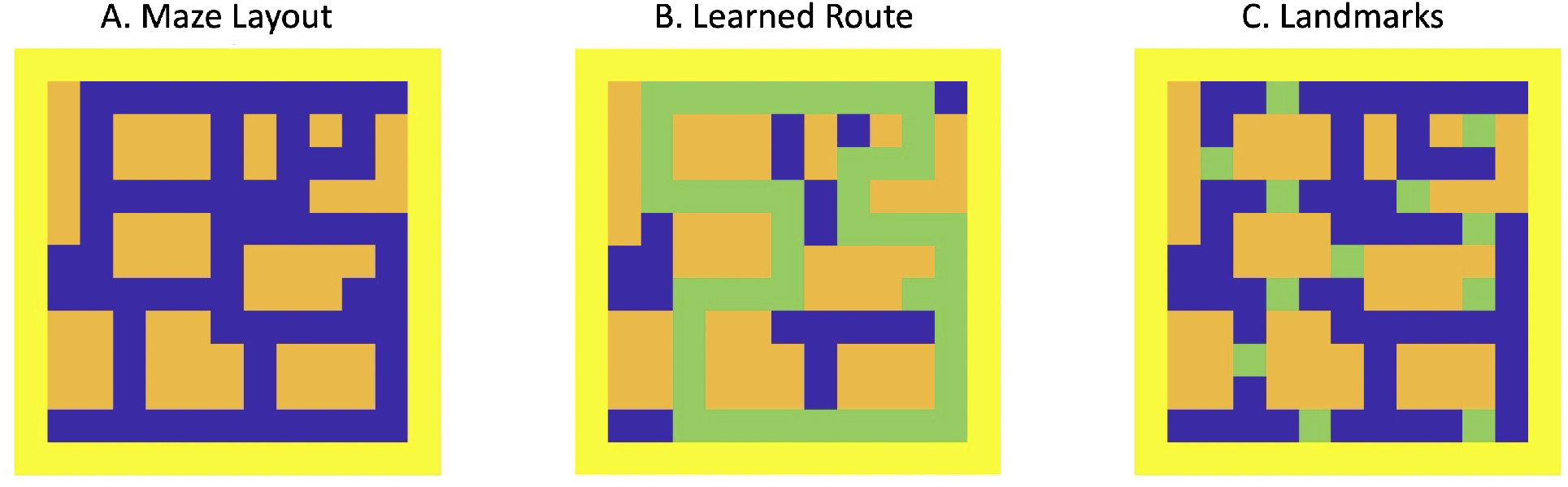
Maps derived from a virtual environment used in human navigation studies [Boone et al., 2019]. A. Map is a 13×13 grid. Yellow denotes the border, orange denotes untraversable areas, and blue denotes traversable areas. B. Green denotes the fixed route learned by participants. C. Green denotes landmark locations used in the testing phase of the experiment.

#### 3.1.1 Experimental Protocol

We converted the virtual maze into a 13×13 grid of Cartesian coordinates, which corresponded to a spiking neural network of the same size. Similar to Boone et al. [2019], we forced the model to take a fixed tour of the environment by presenting the path planning algorithm with starting and ending locations that were close by and along a straight line (see for example, Figure 3 and Figure 13 in Supplemental Materials).

**Figure 3:**
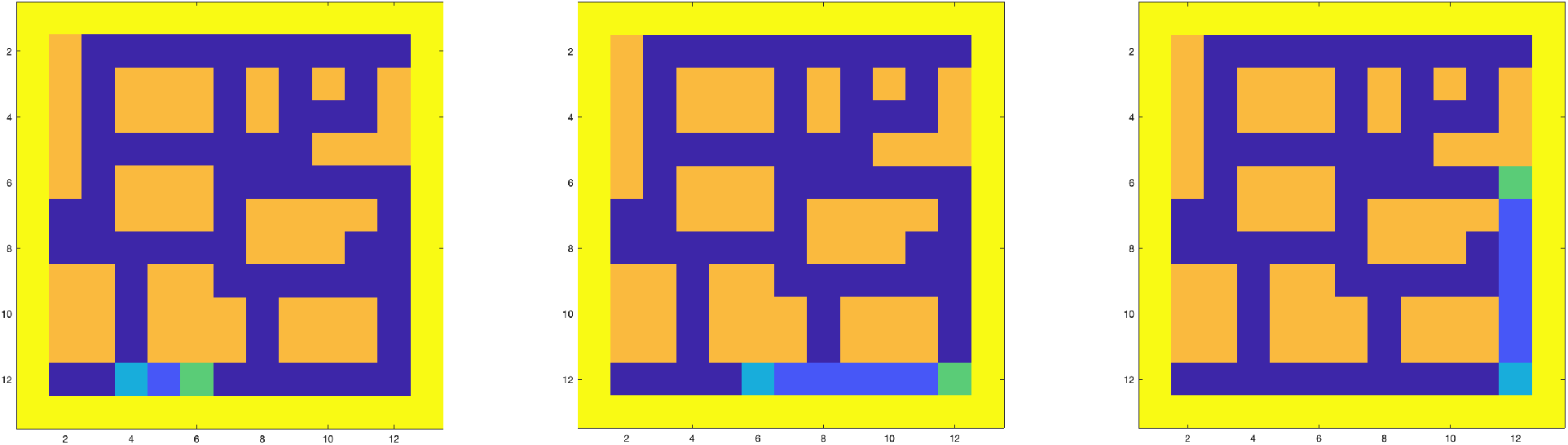
Path calculated by the spike wave algorithm on the first three segments of the learned route. The start is marked in cyan, the goal is marked in green, and the path is shown in light blue for trial 13.

The weights of the spiking neural network, which correspond to axonal delays, were initially set to 10. Through repeated application of E-Prop learning, the weights began to reflect the costs associated with maze features (i.e., traversable regions, boundaries, and obstacles).

#### 3.1.2 Exploration of Model Parameters

The performance of the spike wave propagation algorithm with E-Prop learning is dependent on the amount of training and how long neurons are eligible for updates. We tested the sensitivity of the algorithm by varying the number of training loops and the time constant of the eligibility trace (*τ* in Eqn 6). Similar to the human study, starting and goal locations, which corresponded to landmarks (see Figure 2C), were presented to the algorithm. We chose 24 starting and goal pairs. We measured performance by calculating the number of times the path calculated by the algorithm attempted to navigate over untraversable regions, such as obstacles or outside the maze borders. These errors indicated how well the spiking neural network learned the maze layout.

With some training and a long enough time constant, the neural network learned the maze well enough that there were little or no navigation errors as measured by the number of times a planned path attempted to traverse through an obstacle or boundary. Figure 4 shows the number of errors when the number of training loops and the time constant varied. As the time constant increased greater than 20 timesteps, the number of errors decreased to near zero. The effect of training loops on errors showed a similar trend, but was not as consistent. This is because increasing training on the fixed route does not lead to more exploration of the environment. Based on these exploratory results, we used 6 training loops and a time constant *τ* equal to 25 for the remainder of the simulations in this section.

**Figure 4:**
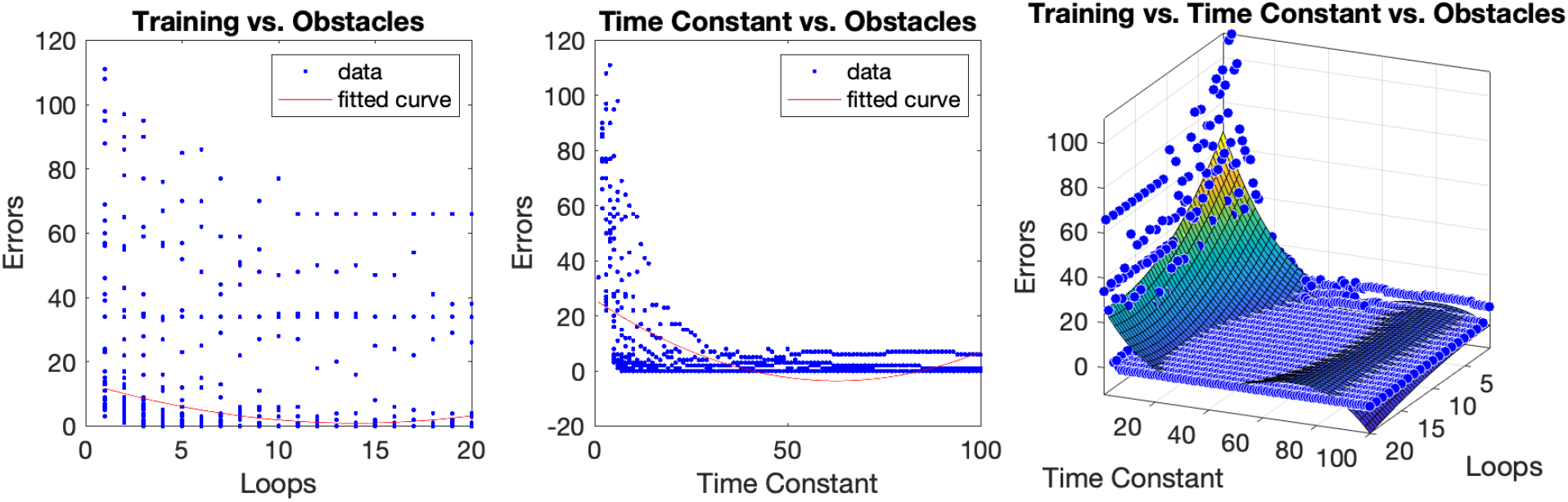
The effect of training and the time constant, *τ*, on errors. Errors are defined as attempts to navigate through obstacles or maze boundaries. Left and middle. Errors as a function of training loops (left chart) and the eligibility trace time constant *τ* (middle chart). Blue dots denote data points. The red line is the fit to the data using a quadratic polynomial curve. Right. Errors as a function of both training loops and the time constant *τ*. Blue dots are data points. The fit to the data was plotted using polynomial surface of degree 2 in x and degree 3 in y.

#### 3.1.3 VTE and Novel Shortcuts

During training on the fixed route, the activity of the spiking neural network reflected the uncertainty of maze features. This could be observed in the number of neurons eligible for updates. For example on the first training loop, more neurons were activated during path planning than on the sixth training loop (see Figure 5). The path planning algorithm will propagate a wave of neural activity based on the weights corresponding to axonal delays. Therefore, early in the training, when features of the maze are unknown, more neurons will be active and eligible (see 6). However, later in the training, the eligible neurons will be confined to the traversable regions in the maze. Most of these eligible neurons will be active on the fixed route. But the spiking wave propagation may activate traversable regions near the fixed route.

**Figure 5:**
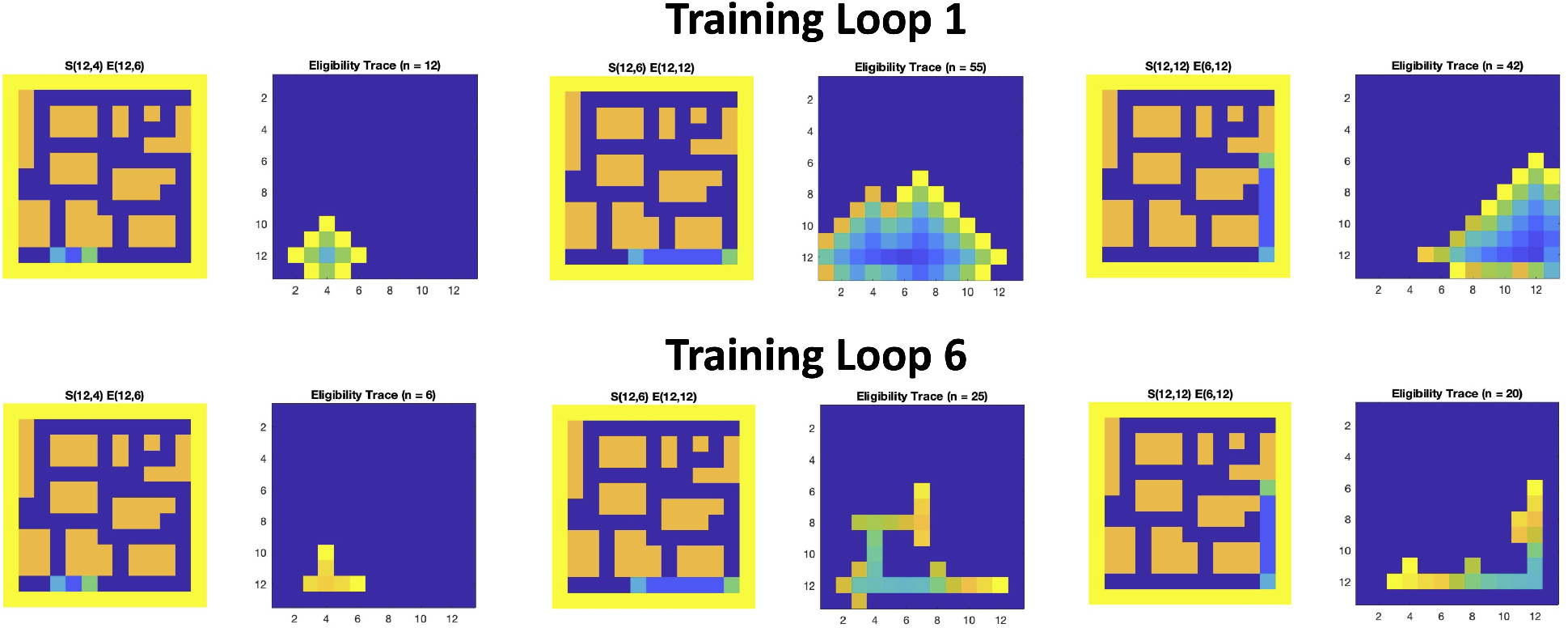
Left panels. Path calculated by the spike wave algorithm for segments of the learned route. The start S(x,y) and goal E(x,y) coordinates are shown above each panel. Right panels. Active neurons during the path planning. The pixels denote neurons with hotter colors corresponding to more recently active neurons according to the eligibility trace. Dark blue pixels denote inactive neurons.

In general, as training progressed, the total loss (*map*_*xy*_ - *D*_*ij*_ in Eqn 5) decreased, which meant the network was learning the maze features, and the number of eligible neurons decreased (see Figure 6). This suggests that as the agent learned its environment, the level of uncertainty and number of VTE episodes decreased.

**Figure 6:**
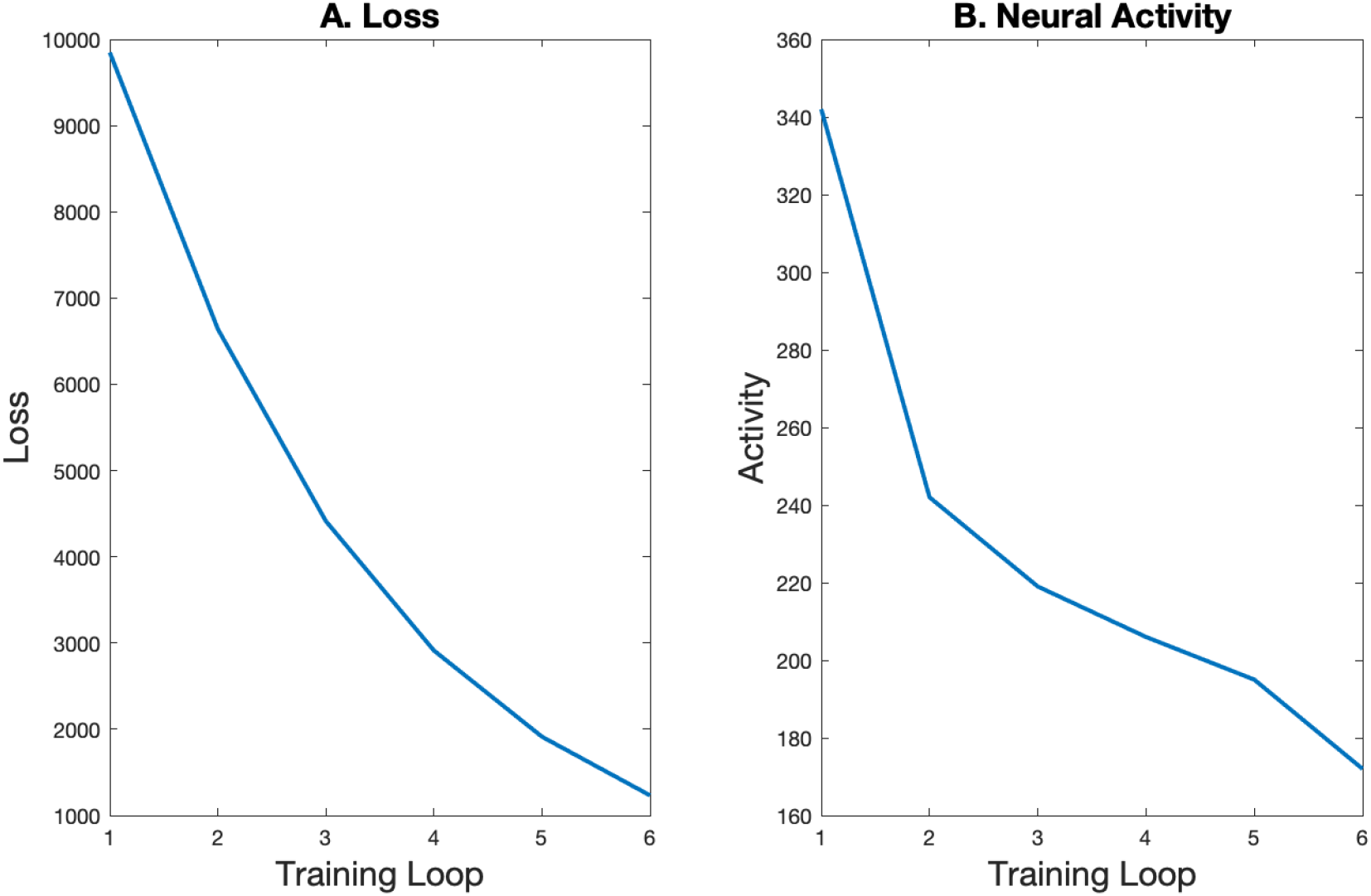
A. Loss per training loop. B. Size of the eligibility trace.

We suggest that the eligibility trace is qualitatively similar to VTE and hippocampal preplay. When the agent was uncertain about the path, there were more neurons active, reflecting a number of alternative routes. This is somewhat like the neural correlates of VTE that have been observed in rodent hippocampus during early exposure to an environment [Johnson and Redish, 2007, Redish, 2016]. After exposure, the neural activity was mostly confined to the route taken. This is somewhat like preplay activity observed in rodent hippocampus when the animal is familiar with an environment ([Redish, 2016, Dragoi and Tonegawa, 2011, Pfeiffer and Foster, 2013].

The neural correlates of VTE may facilitate the construction of cognitive maps and the ability to take novel shortcuts. During training, the eligible neurons spilled over into regions of the environment that were not on the fixed route. This may have led to the calculation of novel routes between landmarks (see Figure 7). The spiking wavefront propagation algorithm calculated novel shortcuts on half of the test trials between landmarks (see Figure 13). Interestingly, adding blockades to the middle section of the fixed route (i.e., coordinates (2,7), (7,7) and (12,7)) led to more shortcuts (15 shortcuts and 9 routes). Similar to Boone et al. [2019], this suggested that there was additional knowledge contained in the neural network, which did not express itself until challenged to do so.

**Figure 7:**
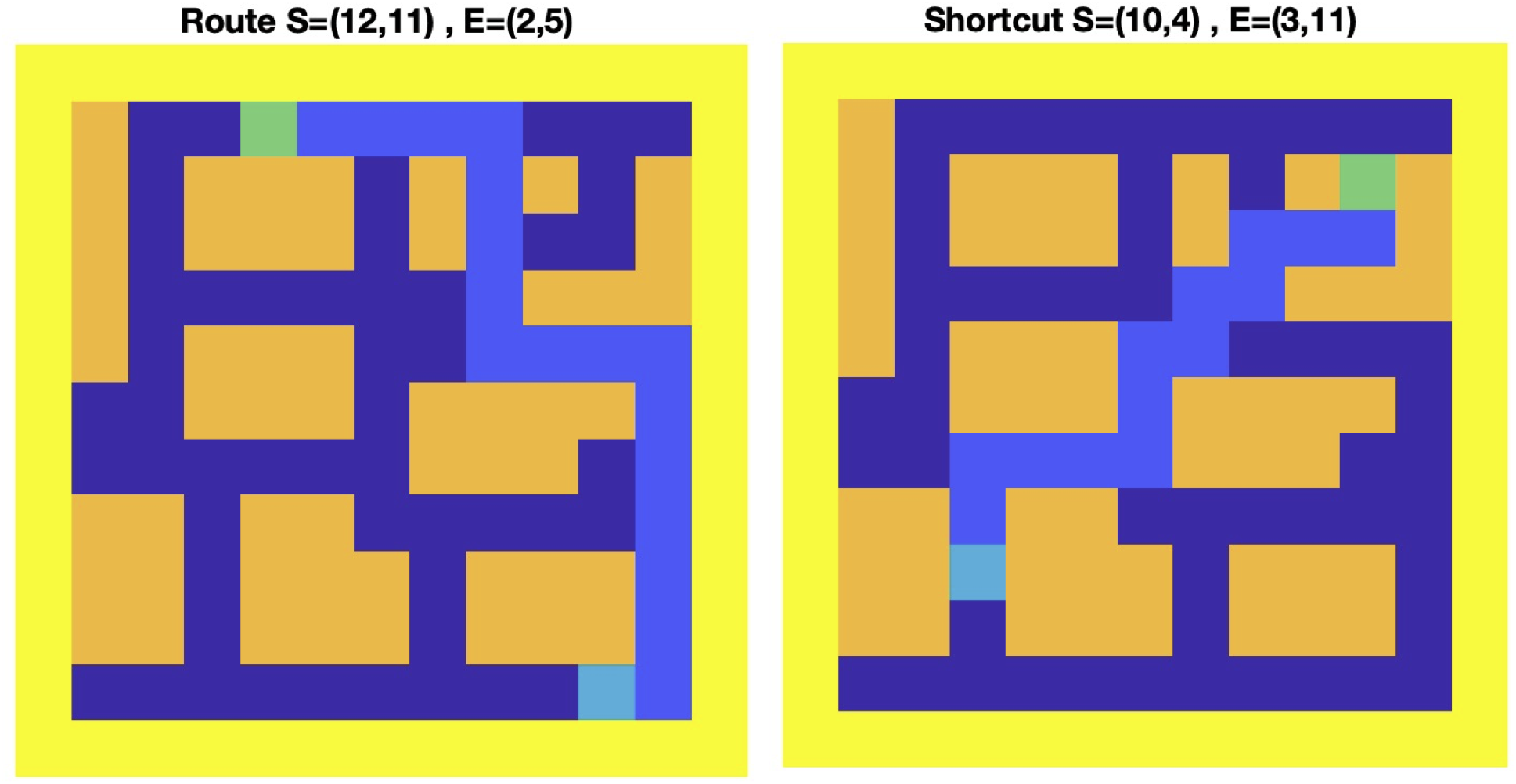
Paths calculated between landmarks. Left. Path along the fixed route. Right. Novel shortcut.

### 3.2 Simulating Rodent Navigation in Tolman Detour Task

In the Tolman detour task, rats were required to choose detours when well-known paths were blocked. Intially, rats were trained to run along a straight corridor from a start location to a goal location (from A to B in Figure 8A). After training, rats needed to plan a detour path when barriers were placed along the original path. If the barrier was placed at P1, the rat could only use the long detour to get from A to B. But if the barrier was placed at P2, the rat might choose either the long detour on the right or the short detour on the left.

**Figure 8:**
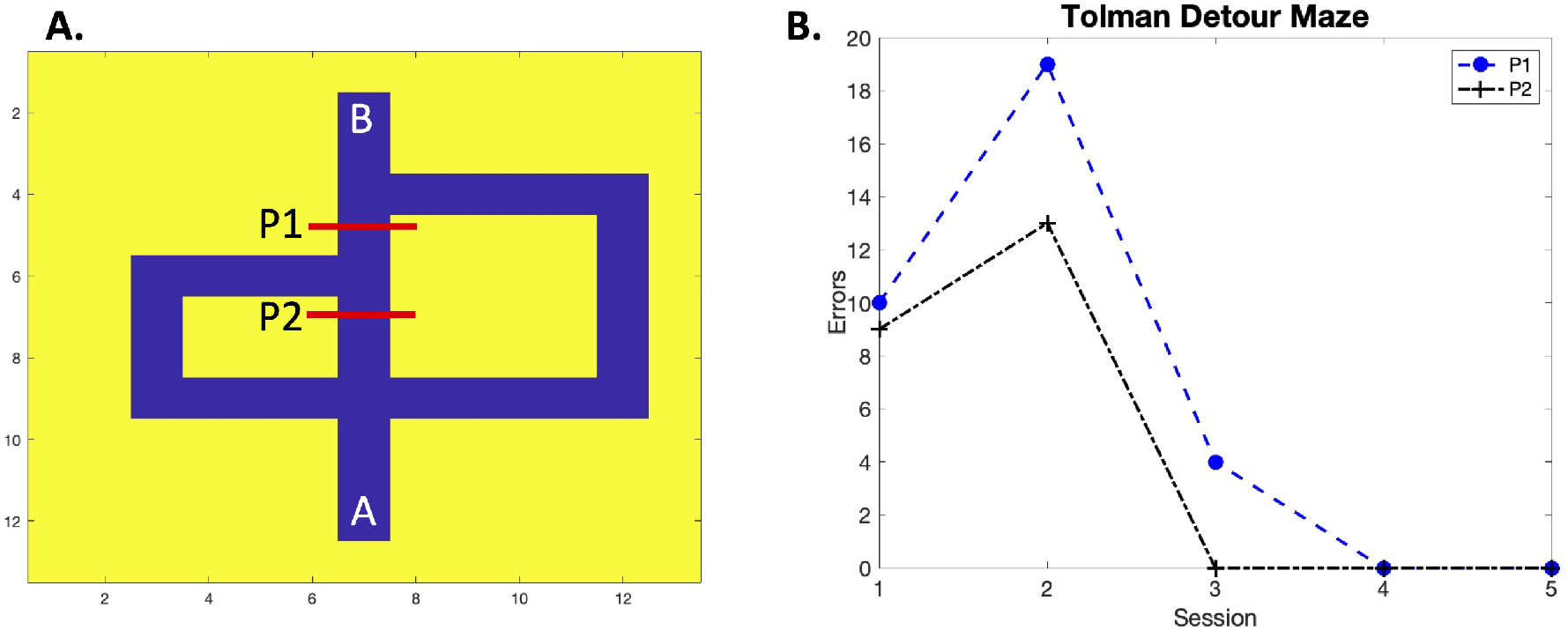
Tolman detour maze task. A. The maze used in the rodent and modeling experiments. The agent is trained to go from A to B. After training, a barrier is placed at P1 or P2. B. Errors during navigation with barriers at P1 or P2. Each session denotes 4 trials from A to B.

#### 3.2.1 Experimental Protocol

We constructed an environment to simulate the Tolman maze. The corridors had a cost of 1 (blue regions in Figure 8A), the maze borders had a cost of 120 (yellow regions in Figure 8A). During the detour test, the barriers, P1 or P2, were placed at (5,7) and (7,7), respectively and the cost was set to 120. The weights in the neural network were initialized to 5.

Initially, the spiking wave propagation algorithm with E-Prop learned a path from A to B over 20 trials. After training, a barrier was placed at P1 or P2. Then the spike wave propagation algorithm underwent 20 trials planning paths from A to B.

#### 3.2.2 VTE and Flexible Rerouting

The spiking wave propagation algorithm with E-Prop quickly learned the best detour given the barrier (see Figure 8B). Each session contained 4 trials where the spike wave propagation planned a path from A to B. Errors were whenever the planned path went outside the corridor or attempted to traverse through a barrier. After the first 2 sessions, the agent planned mostly error-free paths.

Similar to rodent experiments [Alvernhe et al., 2011], when the barrier was placed at P1, the agent took the longer detour (see left panels of Figure 9). Like the rat, the agent attempted to travel straight from A to B when the barrier was introduced. By the 10th trial, the agent took the longer detour to reach its goal. Note how the number of timesteps (TS), the path length (PL), and the loss (see Eqn 5) decreased with the number of trials.

**Figure 9:**
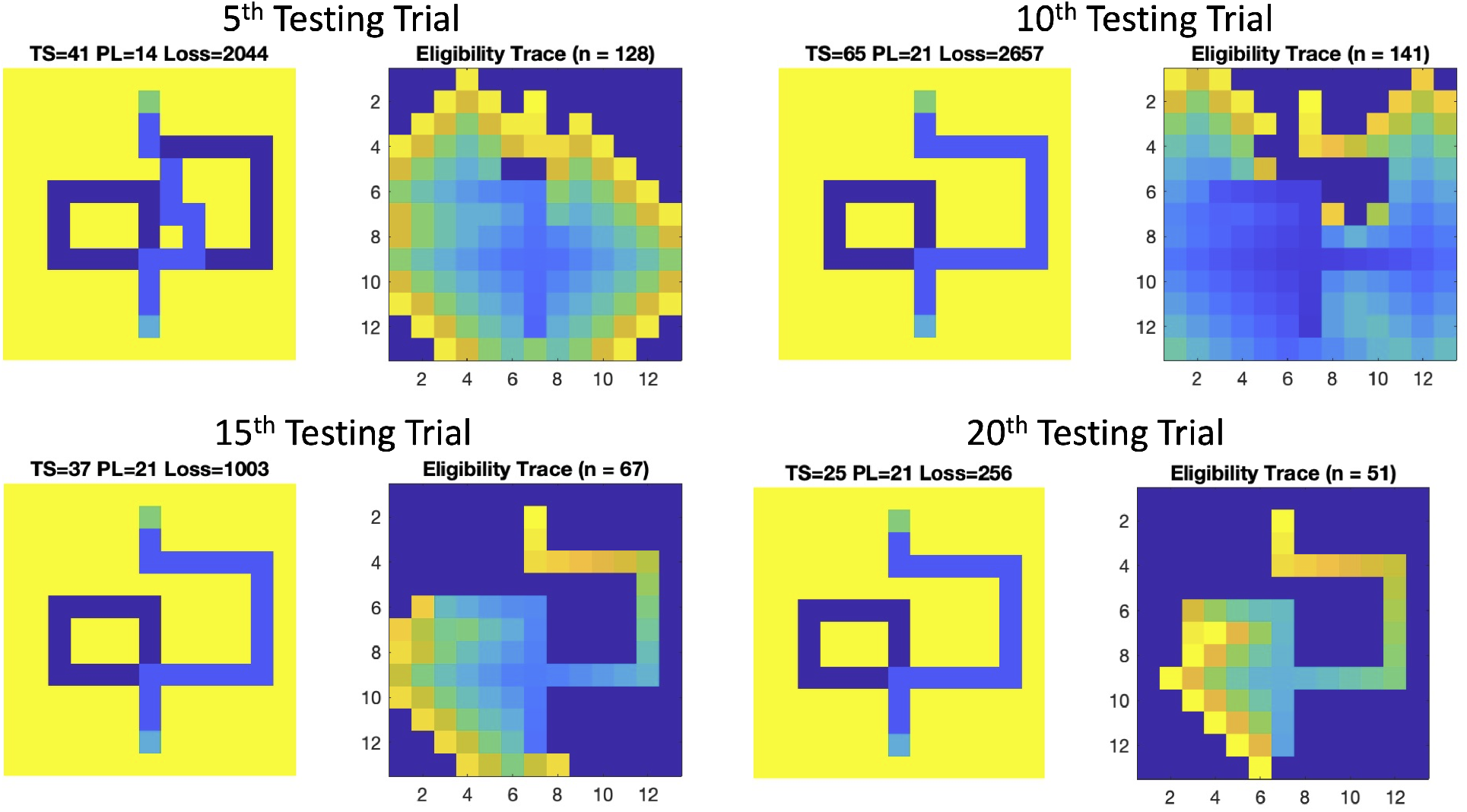
Tolman detour maze task with barrier placed at P1. Left panels. The path planned by the agent. The cyan pixel denotes the start location, and the green pixel denotes the end location. The light blue pixels denote the planned path. The title text contains the timesteps needed to calculate the path (TS), the path length (PL), and the loss when comparing the neural network weights to the maze values. Right panels. The eligible neurons are shown, with the hotter colors signifying more recent activity in the path planning.

When the barrier was placed at P2, the model agent took the shorter detour (see left panels of Figure 10). This differed slightly from the rat experiments. In the rat experiments, the rat occasionally took the long detour, but did favor the short detour. By trial 10, the spiking wave propagation algorithm exclusively planned paths along the shorter detour. Similar to when the barrier was placed at P1, the timesteps (TS), path length (PL), and loss decreased as trials progressed.

**Figure 10:**
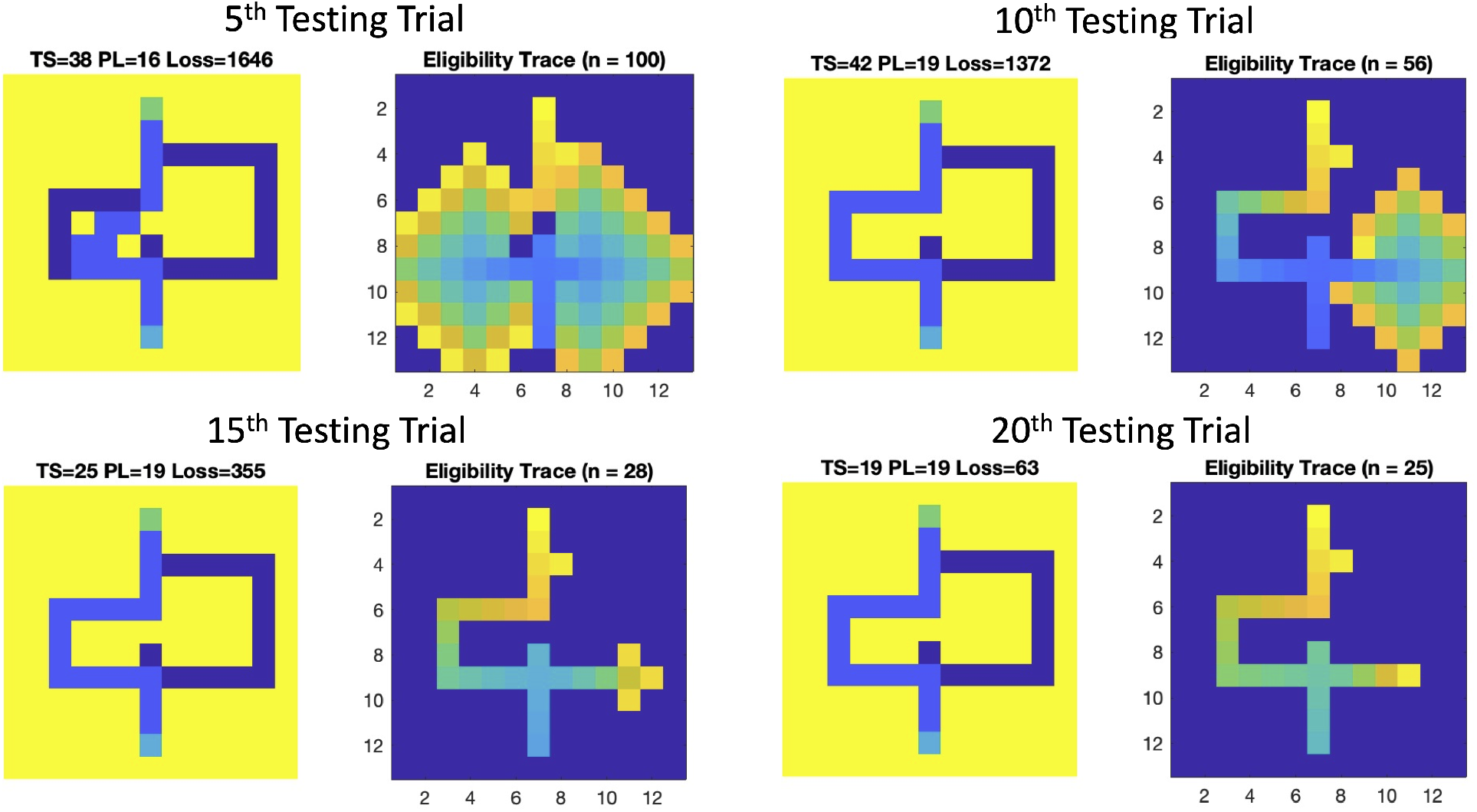
Tolman detour maze task with barrier placed at P2. Notations are the same as in Figure 9.

Introducing a barrier forced the agent to learn alternative routes. This uncertainty was reflected in the eligibility trace of the neural network model (see right panels of Figures 9 and 10). In the first few trials, the model was exploring possible alternatives. We suggest this neural activity is comparable to VTE in the hippocampus. By trial 15 and later, activity was mainly confined to the path taken by the agent. We suggest that this is more akin to preplay in the hippocampus. There are differences however. The activity in later trials still spills over onto alternative paths. Suggesting that these alternatives may be considered. Johnson and Redish [2007] showed neural correlates of VTE, followed by preplay predicting the path on T-mazes. It would be of interest to record the sequence of place cells on a maze such as the Tolman detour maze.

Taken together, these results show how the spiking wave propagation algorithm with E-Prop can rapidly adapt and re-route when changes occur. Furthermore, the uncertainty and consideration of alternative routes can be observed in the neural network activity.

### 3.3 Simulating Rodent Navigation in Morris Water Maze

The Morris water maze is a popular assessment of hippocampal dependent spatial learning in rodents [Morris, 1984]. It is typically conducted in a circular arena filled with opaque water. In one quadrant of the arena, a platform is hidden beneath the surface. In the spatial version of the task, rodents will use distal cues to orient themselves and remember routes to the hidden platform. To prevent the rodent from solving the task with a remembered sequence of movements, the starting position is varied on each trial.

We wanted to test the spiking wave propagation algorithm with E-Prop in a task similar to the Morris water maze. The previous tasks were in mazes that had narrow corridors and sequences of left and right turns. In our version of the Morris water maze, the agent had to navigate an open region where it could move in any direction.

#### 3.3.1 Experimental Protocol

To simulate the Morris water maze, we created a 13×13 map, with an open circular region, which had a diameter or 13, and a platform located at grid location (5,5). The open circular region had a traversal cost that was randomly chosen between 2 and 5. The platform had a cost of 1. All other locations had a cost of 120.

As in the previous simulations, a 13×13 spiking neural network was created with neurons corresponding to the maze grid locations. Since this was an open arena, neurons were connected to their 8 neighbors, which allowed the agent to move in the N, NE, SE, S, SW, W, NW directions. Weights were randomly initialized between 5 and 10. Typically, in the Morris water maze, rats will swim circuitously on early trials before settling into a strategy where they swim directly to the platform from any starting location. In the simulations, random weights and random map costs were chosen to cause the agent to explore its environment, especially during early trials. On each trial, the agent started at a random location on the N, S, E, or W wall. The simulation was run for 32 trials. Ten simulation runs were carried out.

#### 3.3.2 Learning Routes in Open Field

The agent was able to learn routes from any starting location to the platform. Figure 11 shows representative trials from early in the simulation (one of the first 4 trials) and late in the simulation (one of the last 4 trials). In both cases, the spike wavefront propagation algorithm planned paths from the South wall to the platform. Note how in the late trial, the number of timesteps to calculate the path, the path length, and the loss was less than in the early trial. Moreover, the number of eligible also decreased. Because this is an open field, the spike wavefront propagates in all directions. This is because much of the arena is traversable, as compared to the mazes used in our previous simulations. More early and late trials can be seen in Figures 15 and 16 in the Supplemental Materials.

**Figure 11:**
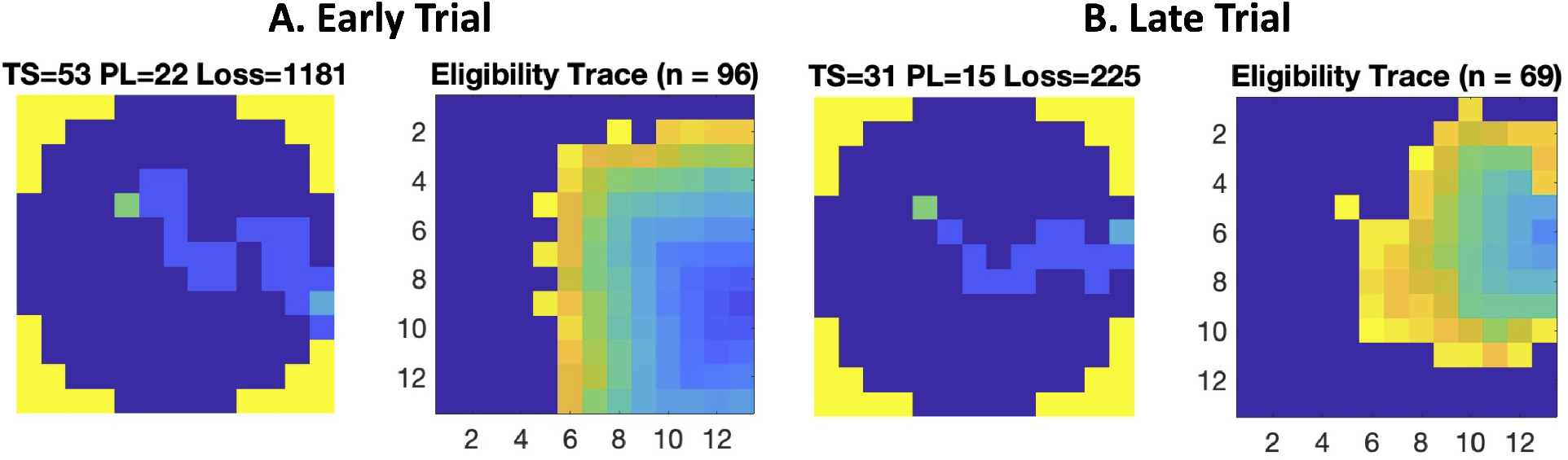
Representative trials from the Morris water maze simulations. Left panels show the path trajectory calculated by the spiking wavefront algorithm with E-prop. The notations in the title are the same as in Figure 9. Right panels show the eligibility of active neurons with warmer colors denoting more recent activity. A. Early trial. B. Late trial.

**Figure 12:**
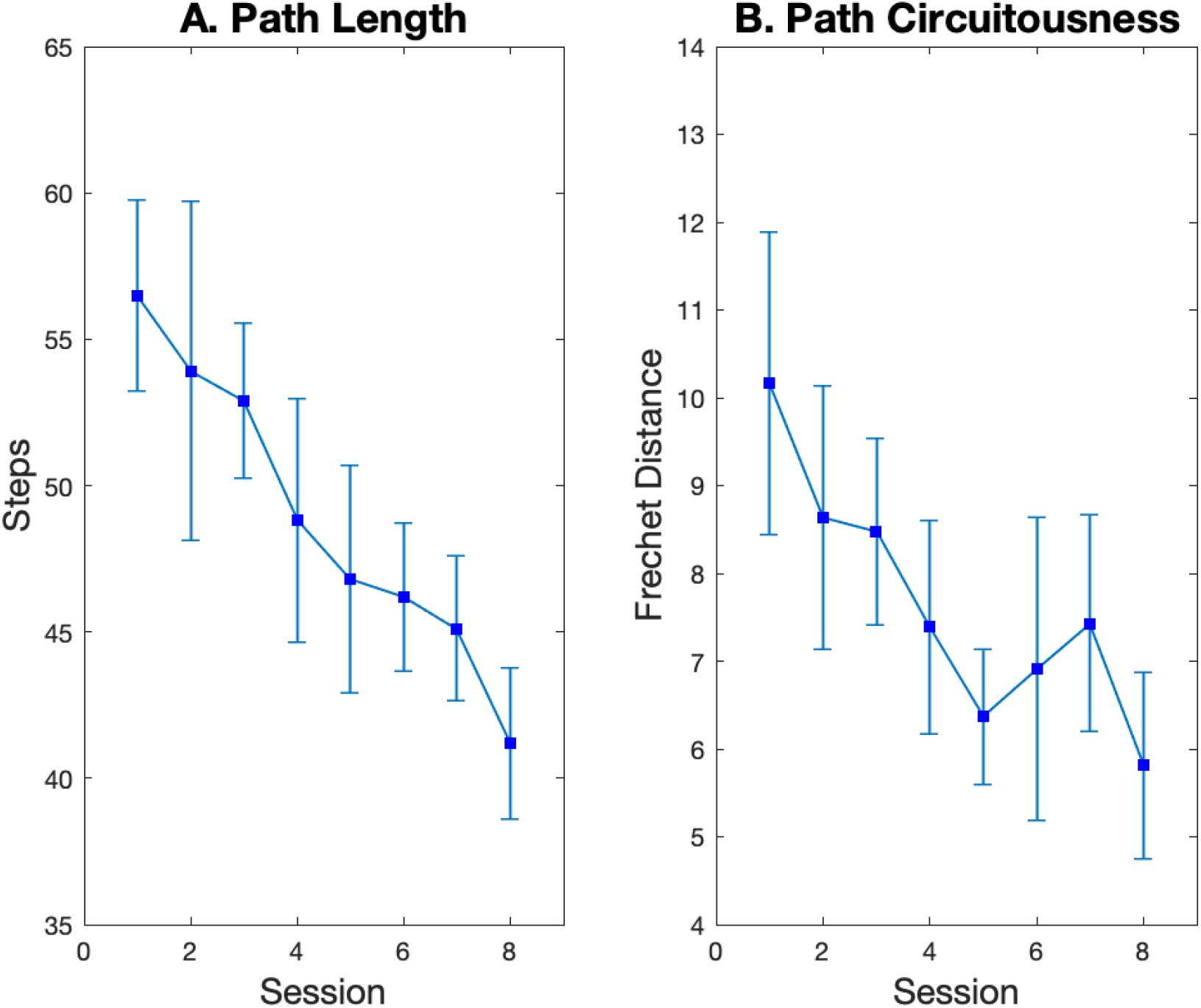
Path statistics for 10 simulation runs of the Morris water maze. There were 8 sessions, each of which contained 4 trials starting from the N, S, E, and W wall. Points denote the mean and the error bars denote the standard deviation. A. Path length. B. Path circuitousness as measured by the Frechet distance Danziger [2020], Eiter and Mannila [1994].

**Figure 13:**
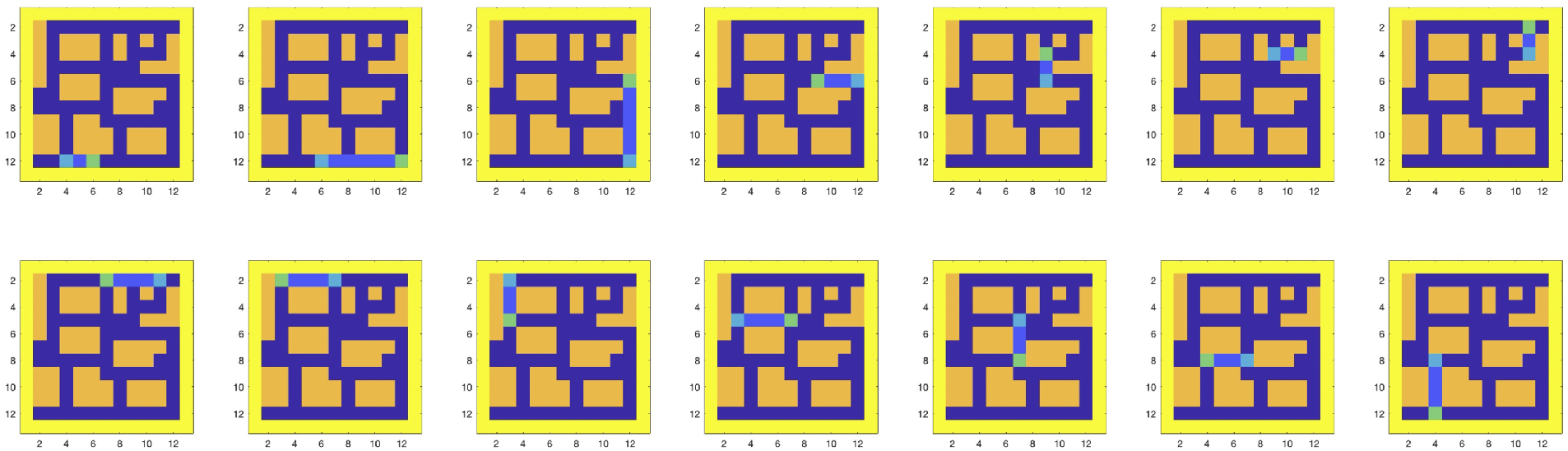
Path calculated by the spike wave algorithm on all segments of the fixed route. The start is marked in cyan, the goal is marked in green, and the path is shown in light blue for trial 13.

**Figure 14:**
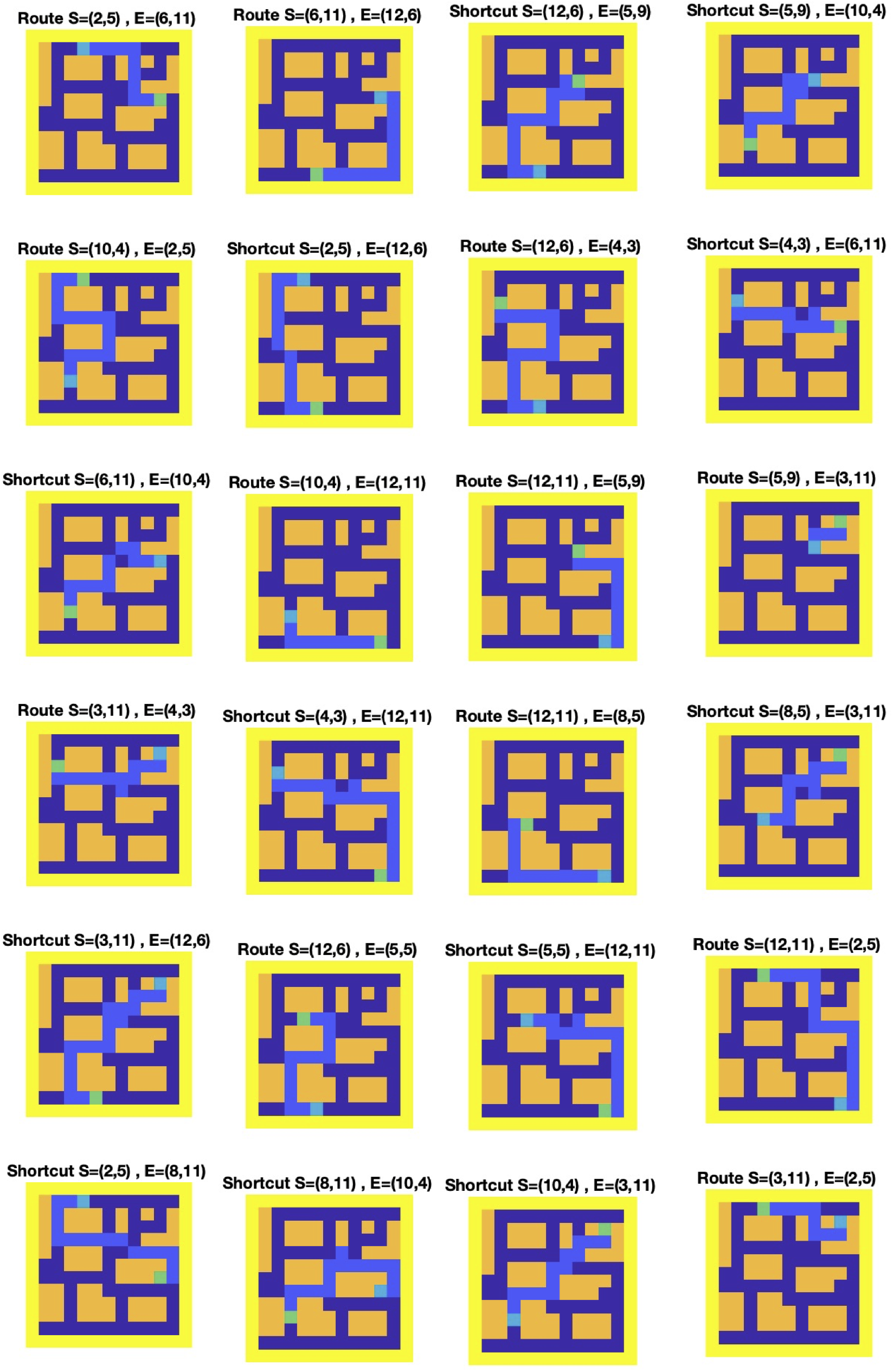
Paths calculated between landmarks for all 24 test trials.

**Figure 15:**
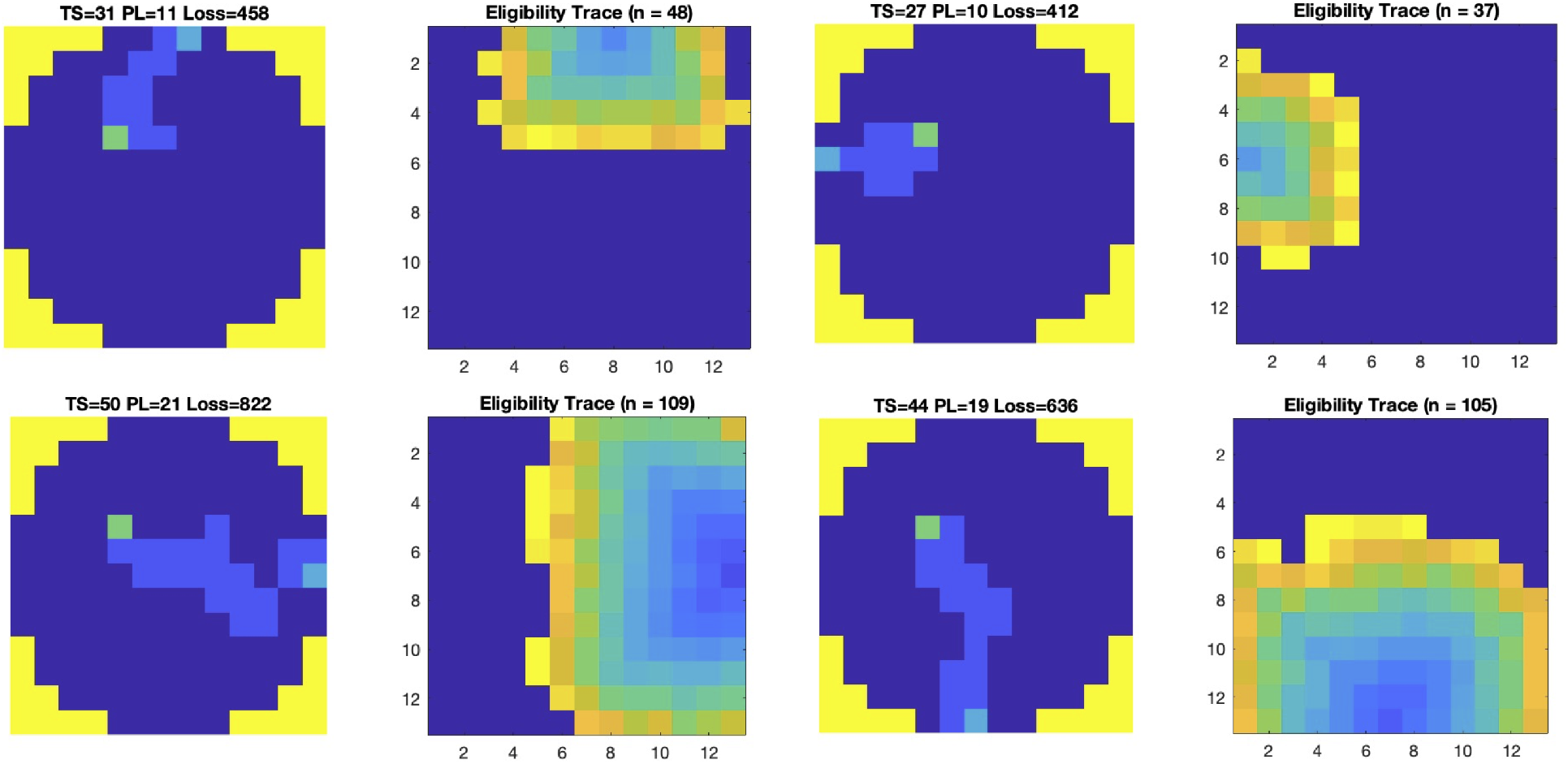
First four trials in Morris water maze simulation.

**Figure 16:**
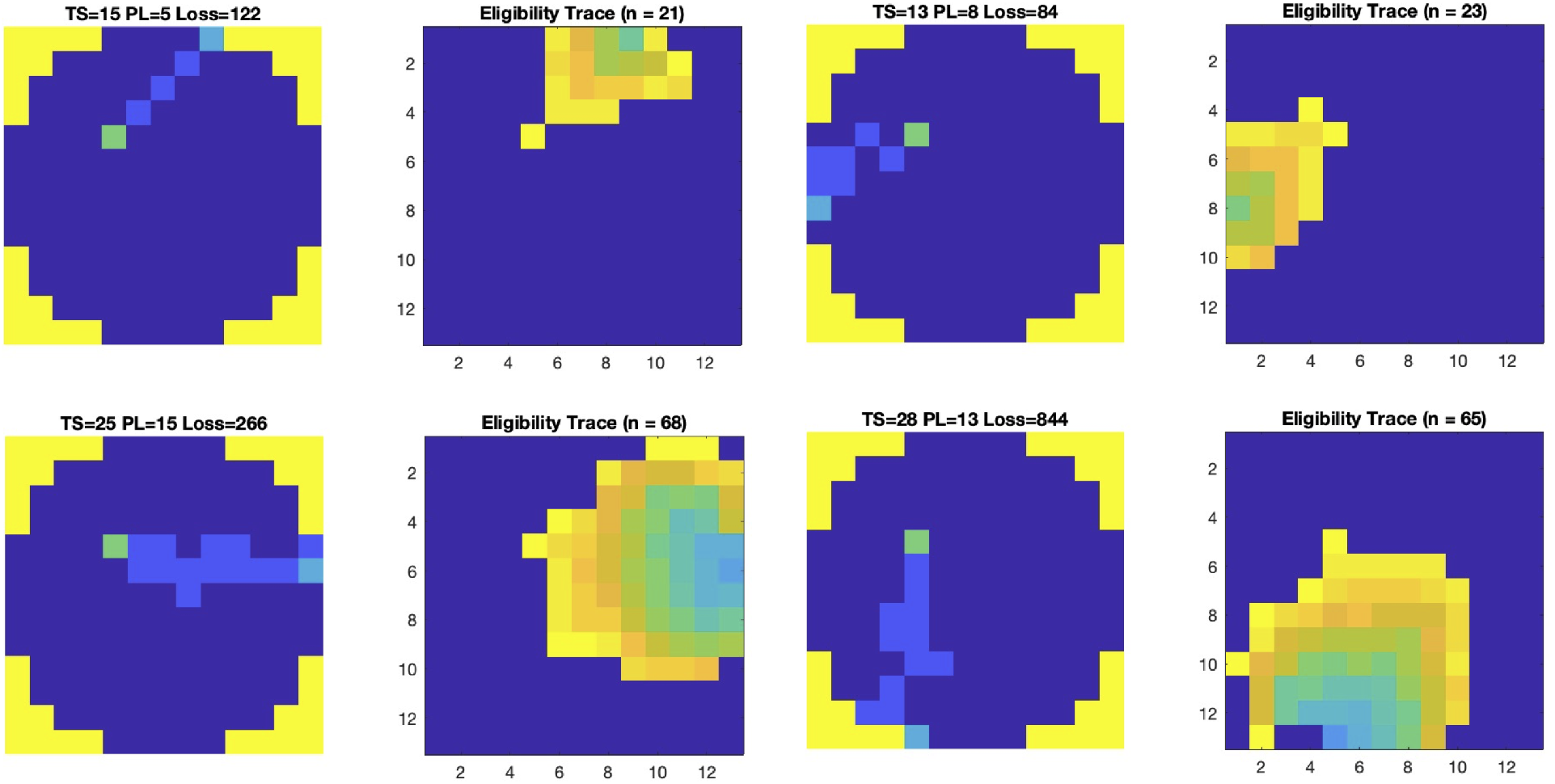
Last four trials in Morris water maze simulation.

As learning proceeded, the paths to the platform became shorter and straighter (see Figure 12). Path circuitousness was measured by calculating the Frechet distance between the spiking wave propagation path and the shortest route from the starting point to the platform [Danziger, 2020, Eiter and Mannila, 1994]. Thus, lower Frechet distances represented less circuitous paths. The agent’s paths differed somewhat from observed rodent swimming. On early trials to the Morris water maze, a rat will swim around the edge of the pool to orient itself. Rats also develop idiosyncratic strategies to solve the maze. To simulate this in our trials, we would have needed to add heuristic strategies to the calculated paths. Still, the overall performance and learning rates are comparable to those observed in the Morris water maze experiments.

## 4 Discussion

A hallmark of cognitive behavior is flexible planning [Miller and Venditto, 2021]. Although the present study and many others concentrate on navigation when investigating flexible planning, such spatial behavior can be extended to non-spatial behavior. In the present study, we show that a path planning algorithm from robotics, coupled with a biologically plausible plasticity rule for learning sequences, could demonstrate flexible path planning in multiple environments, and generate neural correlates of assessing alternatives when faced with uncertainty. Our simulations suggest that Vicarious Trial and Error or VTE is not only a neural correlate of uncertainty during decision-making, but also leads to information gathering for future planning.

### 4.1 Path Planning Algorithms

Planning routes and paths are behaviors that both animals and robots must perform under environmental constraints, such as limited energy and changing conditions. Path planning has commonly been applied to robot applications [LaValle, 2011a,b]. Dijkstra’s algorithm is a well-known approach for finding the shortest paths. The A^*^ path planning algorithm is a faster variant of Dijkstra’s algorithm that selects the next nodes to explore using some heuristic cost function. Similar to Dijkstra and A^*^, wavefront planners, potential fields, and diffusion algorithms can calculate optimal paths [Barraquand et al., 1991, Soulignac, 2011]. However, all of these algorithms can be computationally expensive if applied to large-scale maps.

The spiking wavefront propagation path planner presented here has been used for robot navigation [Hwu et al., 2018], and to simulate human navigation [Krichmar and He, 2021]. Unlike other path planners, the spiking wavefront algorithm is a network of spiking neurons. Therefore, the algorithm is compatible with power efficient neuromorphic hardware, as was shown by implementing it on the IBM TrueNorth NS1e [Fischl et al., 2017]. The spiking wavefront propagation algorithm is also supported by the observation of spreading activation of hippocampal place activity prior to taking action [Dragoi and Tonegawa, 2011, Pfeiffer and Foster, 2013]. This preplay of activity may have correlates to path planning.

The present work explores how a biologically plausible learning rule [Bellec et al., 2020], specifically designed for spiking neural networks, could extend the spiking wavefront propagation algorithm. Rather than having learning be related to changes in synaptic efficacy, the present algorithm changed the delays between pre- and post-synaptic neurons This was inspired by evidence suggesting that the myelin sheath, which wraps around and insulates axons, undergoes a form of activity-dependent plasticity [Fields, 2015]. These studies have shown that the myelin sheath becomes thicker with learning motor skills and cognitive tasks. A thicker myelin sheath implies faster conduction velocities and improved synchrony between neurons. Although, we are not suggesting that hippocampal learning is solely due to these changes in axon propagation, it is an intriguing form of plasticity that is rarely applied to neural networks.

### 4.2 Comparison to Other Models

In addition to path planning algorithms for robotics, there have been a number of neurobiologically inspired models of path planning. Many of these models contain simulated hippocampal place cells and entorhinal grid cells. For example, it has been proposed that the place cells form a topological navigation system and the grid cells form a vector navigation system [Edvardsen et al., 2020]. When the agent encounters an obstacle, the planner re-routes through hippocampal replay. Other models suggest that interaction between the hippocampus and the neocortex can support vector-based navigation. Models of the interplay between the prefrontal cortex and a set of hippocampal place cells have shown the ability to plan novel shortcuts and paths [Cazin et al., 2019, Martinet et al., 2011]. Vector navigation, which supports planning shortcuts, has been shown in deep neural networks that have a grid code [Banino et al., 2018]. These models suggest that grid cells are important for planning direct trajectories to goals.

Some form of preplay appears to be important for flexible future planning. For example, a network model of head direction cells, grid cells, place cells, and prefrontal cortex cells found shortcuts in the Tolman detour maze [Erdem and Hasselmo, 2012]. The grid cells drove place cells along look-ahead trajectory similar to hippocampal preplay. The Tolman detour maze was also solved using a Successor Representation model [Stachenfeld et al., 2017].

Although these models demonstrate forward planning and make predictions regarding hippocampal forward replay or preplay, as well as the function of entorhinal grid cells, they do not address the uncertainty and weighing of alternatives when the environment is novel or perturbed. VTE was originally suggested by Tolman and his colleagues when they observed the rat moving its head back and forth to assess different alternatives [Tolman, 1948]. Later, it was observed that these head movements corresponded to sweeps of hippocampal place activity along these alternatives [Johnson and Redish, 2007, Redish, 2016].

#### 4.2.1 VTE facilitates information gathering for future planning

The spiking wavefront propagation path planner presented here has activity similar to the neural correlates of VTE: 1) It is more prominent during early learning and when faced with environmental challenges. 2) The wave of activity propagates to alternative choices, and 3) After experience, the wave of activity more resembles preplay in that it becomes confined to the future choice.

Our simulations suggest another function of VTE, which is information gathering for future use. In the present algorithm, the activity propagated to portions of the maze that were not visited. Nevertheless, the E-Prop learning with BPTT stored this information in the network weights. So when a novel shortcut was available, as in the simulation of human navigation, or when a new route was necessary, as in the Tolman detour task, the agent was able to express this stored information and rapidly adapt and re-route.

Unlike observations in the rodent, the present model’s VTE propagates in parallel, rather than sequential [Johnson and Redish, 2007, Redish, 2016]. However, it should be noted that this occurs with the neural network having no knowledge of the maze structure. It may be that rats in a corridor maze know that paths are constrained by the maze walls. Another difference is the movement of the rat’s head coincides with the activation of a VTE. This sequential trace might be triggered by the rat’s visual system. It would interesting to test the present model on a robot with active vision to see if sequential VTE-like activity emerges when the robot looks in a particular direction. The performance of the spiking wavefront propagation algorithm on the Morris water maze predicts that goal-driven behavior in open environments might demonstrate VTE’s that are more diffuse. Pfeiffer and Foster [2013] showed sequential preplay in an open-field arena, but did not record neural activity when the arena was unfamiliar to the rat. It would be of interest to see if the diffuse VTE-like activity presented here resembles CA1 activity on early trials in an open field environment.

## 5 Conclusion

We propose a novel model of VTE and preplay based on a path planning algorithm used in robot navigation. The introduction of a biologically plausible learning rule allows the model to simultaneously learn its environment and plan paths. In simulations of human and rodent navigation, it was observed that our algorithm could flexibly plan efficient paths and rapidly recover from perturbations.

## 6 Acknowledgments

This material is based upon work supported by the United States Air Force Research Laboratory (AFRL) and Defense Advanced Research Projects Agency (DARPA) under Contract No. FA8750-18-C-0103. Any opinions, findings and conclusions or recommendations expressed in this material are those of the author(s) and do not necessarily reflect the views of the United States Air Force Research Laboratory (AFRL) and Defense Advanced Research Projects Agency (DARPA).

